# Structural and biochemical insights into lipid transport by VPS13 proteins

**DOI:** 10.1101/2022.03.11.484024

**Authors:** Jyoti Adlakha, Zhouping Hong, PeiQi Li, Karin M. Reinisch

**Author notes:** Aligning Science Across Parkinson’s (ASAP) Collaborative Research Network, Chevy Chase, MD, 20815, USA. these authors contributed equally (Jyoti Adlakha, Zhouping Hong).

## Abstract

VPS13 proteins are proposed to function at contact sites between organelles as bridges for lipids to move directionally and in bulk between organellar membranes. VPS13s are anchored between membranes via interactions with receptors, including both peripheral or integral membrane proteins. Here we present the crystal structure of VPS13s adaptor binding domain (VAB) complexed with a Pro-X-Pro peptide recognition motif present in one such receptor, the integral membrane protein Mcp1p, and show biochemically that other Pro-X-Pro motifs bind the VAB in the same site. We further demonstrate that Mcp1p and another integral membrane protein that interacts directly with human VPS13A, XK, are scramblases. This finding supports an emerging paradigm of a partnership between bulk lipid transport proteins and scramblases. Scramblases can re-equilibrate lipids between membrane leaflets as lipids are removed from or inserted into, respectively, the cytosolic leaflet of donor and acceptor organelles in the course of protein-mediated transport.

---

VPS13 family proteins are thought to mediate directional bulk glycerolipid transfer between organelles at contact sites, where organelles are closely apposed, effecting membrane expansion and organelle biogenesis. These proteins feature a single extensive beta sheet that is highly curved to form a taco-shell like structure whose concave surface is lined entirely with hydrophobic residues. The taco-shell is long enough to span the cytosolic space between organelles, and the hydrophobic groove along its length can solubilize tens of lipid fatty acid moieties. Recent and ongoing studies are identifying elements at both ends of the proteins that interact with organellar membranes either directly or via adaptor proteins to position the taco-shell as a bridge between organelles, allowing lipids to travel between membranes along the hydrophobic groove (reviewed in (Dziurdzik and Conibear, 2021; Leonzino et al., 2021)). So far the best characterized members of the family are VPS13 itself, including a single homolog in fungi and four in humans (A-D), and ATG2, whose lipid transfer function is required for autophagosome biogenesis. The human VPS13 proteins are of significant biomedical interest because their dysfunction is linked to severe neurological diseases, including Parkinson’s (VPS13C)(Lesage et al., 2016) and chorea acanthocytosis (VPS13A)(Rampoldi et al., 2001; Ueno et al., 2001). Among the VPS13 proteins, *S. cerevisiae* Vps13p has been most thoroughly studied and has served as a model system to understand VPS13 function in humans (Dziurdzik et al., 2020; Park et al., 2021).

A major question for VPS13 family proteins is how they localize to and associate with organellar membranes to allow for efficient lipid transfer between the transfer protein and membranes. In yeast, Vps13p localizes to multiple contact sites in part via the so-called “VPS13 adaptor binding domain”, or VAB, near the C-terminal end (Figure 1A)(Bean et al., 2018). The VAB, with no significant sequence similarity to any previously characterized protein domain, interacts with a Proline-X-Proline motif present in receptor proteins at contact sites (Bean et al., 2018). Receptor proteins include the sorting nexin Ypt35p on the endosome, Spo71p on the prospore membrane, and a multi-pass integral membrane protein, Mcp1p, in the outer mitochondrial membrane (Bean et al., 2018; John Peter et al., 2017). The VAB is present in all VPS13s (but not other VPS13 family proteins like ATG2), including human VPS13 A-D, and likely plays a similar role in their localization, though its human interactome is still unknown and may feature different recognition motifs other than Pro-X-Pro. Here we present the crystal structure, at 3.0 Å resolution, of the VAB from the fungus *Chaetomium thermophilum*, complexed to the Pro-X-Pro motif of Mcp1p from the same organism. We also present biochemical confirmation that the Pro-X-Pro-motifs from Ypt35p and Spo71p bind at the same site as the Mcp1p Pro-X-Pro motif, in agreement with competition studies (Bean et al., 2018).

**Figure 1.**
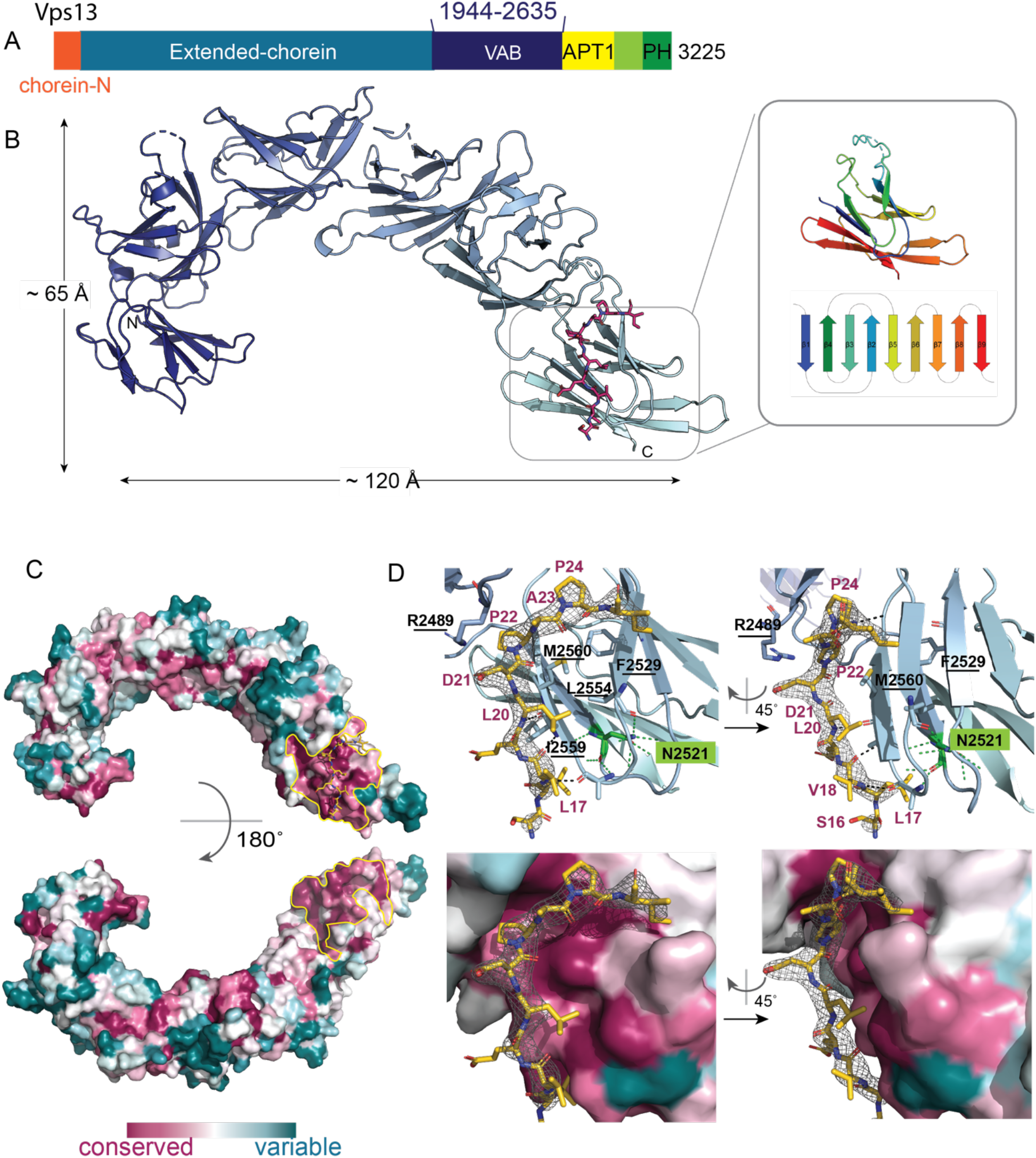
Architecture of the VAB and its Pro-X-Pro motif binding site. (A) Schematic showing domain architecture of Vps13p. Residue numbers refer to the *Chaetomium thermophilu*m Vps13p sequence. (B) Ribbons diagram for the VAB from *C. thermophilum*, showing its 6 repeated modules. The Pro-X-Pro motif is colored magenta. Inset shows one module, colored from blue at the N-terminus to red at the C-terminus, and a topology diagram colored in the same way. Supplementary Figure 1A shows differences between the crystal structure and the AlphaFold2 prediction. (C) Sequence conservation, based on an alignment of 1000 fungal Vps13s as determined by Consurf (Ashkenazy et al., 2016) and mapped onto the VAB surface. A patch centered on the 6^th^ repeat and including the interface between the 5^th^ and 6^th^ repeats, outlined in yellow, is highly conserved and is the binding site for the receptor Pro-X-Pro motif. For electrostatic potential mapping, see Supplementary Figure 1B. (D) Difference density from a 2Fo-Fc map (3.0 I/sigma contour level), into which the Pro-X-Pro binding motif was modeled, is shown. Top: Two views of the Pro-X-Pro binding motif (yellow) bound to the VAB (light blue). Residues in the VAB binding surface, including those that were mutated to abrogate binding of the Pro-X-Pro motif, are labeled (mutated residues underlined). The asparagine in the 6^th^ module at the end of ß1, which is conserved in all modules, and which was mutated in previous interaction studies (Dziurdzik et al., 2020), is labeled (N2521). In the protein interior, it is part of an extensive hydrogen bonding network that stabilizes module folding (Supplementary figure 1C).

Parallel studies with the VPS13-family protein ATG2 indicated that ATG2 localizes to contact sites between the ER, where most lipids are synthesized, and nascent autophagosomes, and that ATG2 interacts with scramblases both in the ER and in the autophagosome (Ghanbarpour et al., 2021; Maeda et al., 2020; Matoba et al., 2020). We and others proposed previously that the scramblases allow for directional bulk lipid flow by restoring the bilayer symmetry of both the donor and acceptor membranes as lipids are asymmetrically removed from or inserted into their cytosolic leaflet in the course of protein-mediated lipid transfer. Here we present evidence that the VPS13s themselves also work with lipid scramblases, showing scramblase activity for both Mcp1p, the Pro-X-Pro motif containing protein that interacts with yeast Vps13p at mitochondria (Bean et al., 2018; John Peter et al., 2017), and for XK (also known as XKR1), which interacts with VPS13A in humans (Park and Neiman, 2020). XK-related proteins −4, −8, and −9 are known caspase-activated scramblases in the plasma membrane (Suzuki et al., 2014), but no such activity has been previously reported for XK itself. Our data support a model in which bulk lipid transporters in the VPS13 family function in collaboration with scramblases.

## Results

To better understand how VPS13s might interact with organellar membranes, we undertook a structural characterization of the VAB. While we were unsuccessful in crystallizing the VAB domain by itself, we did obtain crystals of a construct comprising the VAB of *C. thermophilum* Vps13p (residues 1944-2635) N-terminally fused to the Pro-X-Pro-motif of Mcp1 (residues 15-32), PXP(Mcp1_ct_)-VAB_ct_. The crystals belong to spacegroup P2_1_and diffract to 3.0 Å. We used molecular replacement for phasing with a model for the VAB derived from Alphafold2 (Jumper et al., 2021). The VAB comprises six modules of the same topology arranged end-to-end to resemble a fish hook (Figure 1B). Each module features nine beta-strands connected by loops and arranged into a beta-sandwich, similar to that of lysins such as ostreolysin (PDBID 6MYI) or hemolysin (PDBID 6ZC1) (Figure 1B, inset). There are two VAB molecules in each asymmetric unit. We were able to model all six of the modules in one copy, PXP-VAB_1-6_. In the second copy, PXP-VAB_1-5_, we modeled only the first five repeated modules (residues 1944 – 2516) as the electron density for the sixth repeat was poor, most likely because this repeat is flexibly positioned with respect to the rest of the protein. The first five modules for the two copies of the VAB in the crystal superimpose closely (rmsd over 428 Cα’s is 0.48 Å). (A comparison of PXP-VAB_1-6_ and the AlphaFold2 prediction are shown in Supplementary Figure 1A; major differences exist at the interfaces between repeated modules.) One face of *C. thermophilum* VAB is largely acidic whereas the other has a large basic patch (Supplementary Figure 1B). A prominent patch on the surface of the 6^th^ module and extending to the interface between the 5^th^ and 6^th^ modules is highly conserved across fungi, indicating functional importance (Figure 1C). Indeed, this surface includes the binding site for the Pro-X-Pro motif (below). A similar surface, on the 6^th^ repeat and at its interface with the 5^th^ repeat, is highly conserved in comparisons of metazoan VPS13 VABs as well, indicating a key function, likely also in receptor binding. The patch residues are less well conserved in comparing fungal and metazoan VABs, however, possibly because the metazoan VPS13s recognize different receptor motifs, other than Pro-X-Pro.

There is strong density for the Pro-X-Pro-motif fused to the N-terminus of PXP-VAB_1-5_ (L_17_V_18_E_19_L_20_D_21_**P**_22_A_23_**P**_24_I_25_A_26_E_27_), showing that it interacts in trans with PXP-VAB_1-6_, binding on the acidic face of the VAB at the conserved interface between its fifth and sixth repeats (Figure 1D). Residues L17-D21 form a short ß-strand along side and parallel to the fourth ß-strand of the 6^th^ repeat module, with the two conserved hydrophobic residues (L17 and L20) packed against hydrophobic surfaces. The two proline side chains (P22, P24) are buried in a groove that runs along the interface. The strong conservation of residues forming the Pro-X-Pro-binding site in PXP-VAB_1-6_ support that this binding surface is physiologically relevant and not an artifact of crystallization. The Pro-X-Pro motif at the N-terminus of PXP-VAB_1-6_ appears to be disordered, as there is no density for it in the electron density maps. Likely, because of crystal packing constraints, the peptide cannot access the binding site in PXP-VAB_1-5_, or the binding site is not formed in PXP-VAB_1-5_ (the 6th module is not visible in the density), or both.

A Pro-X-Pro binding site at the interface of the fifth and sixth modules of the VAB is consistent with previously reported biochemical experiments, indicating that the fifth and sixth repeat modules are sufficient for binding the Pro-X-Pro motif (Dziurdzik et al., 2020). Further, mutation of an asparagine residue at the C-terminal end of first ß-strand of the sixth repeat module of the VAB (N2521 in *C. thermophilum* Vps13p) abolished Pro-X-Pro binding (Dziurdzik et al., 2020). This asparagine, in the protein interior, does not form part of the Pro-X-Pro motif binding surface of the VAB as observed in the crystal structure (see Figure 1E), but is highly conserved in all modules, as part of an extensive hydrogen bonding network stabilizing the module’s core (including the ß3- ß4 hairpin at the Pro-X-Pro peptide binding site, Supplementary Figure 1C). Its mutation to an alanine very likely destabilizes the sixth repeat module, at least locally, preventing formation of the Pro-X-Pro binding site. (Mutation of the corresponding asparagine in the most N-terminal repeat module, which is far removed from the identified Pro-X-Pro binding site, was also reported to abolish VPS13 interactions with its receptor proteins. Possibly this mutation causes local destabilization to perturb packing of the first against the second VAB repeat modules, thus somehow affecting the structure of the VAB or its positioning within the full-length VPS13 to interfere with receptor binding.)

Further characterization of the *C. thermophilum* VAB alone, the PXP(Mcp1_*ct*_)-VAB_*ct*_ construct used in crystallization, and a construct in which the Pro-X-Pro-motif is C-terminally fused to the VAB, VAB_ct_-PXP(Mcp1_ct_), further support the proposed binding site between the fifth and sixth modules. Both size exclusion chromatography and negative stain electron microscopy indicate that the VAB alone and VAB_ct_-PXP(Mcp1_ct_), where the Pro-X-Pro-motif can access the proposed binding site in cis, are monomeric (Figure 2 A-D). In contrast, PXP(Mcp1_*ct*_)-VAB_*ct*_ forms dimers (Figure 2 A,D). In this case, the N-terminal Pro-X-Pro-motif is too far from the binding site to access it in cis and instead accesses it in trans (Figure 2B). When we introduced mutations to perturb the identified binding site (R2489E, L2554A, I2559A and M2560A; see Figure 1D), PXP(Mcp1_*ct*_)-VAB_*ct*_ no longer dimerized, indicating that the Pro-X-Pro-motif no longer binds (Figure 2D).

**Figure 2.**
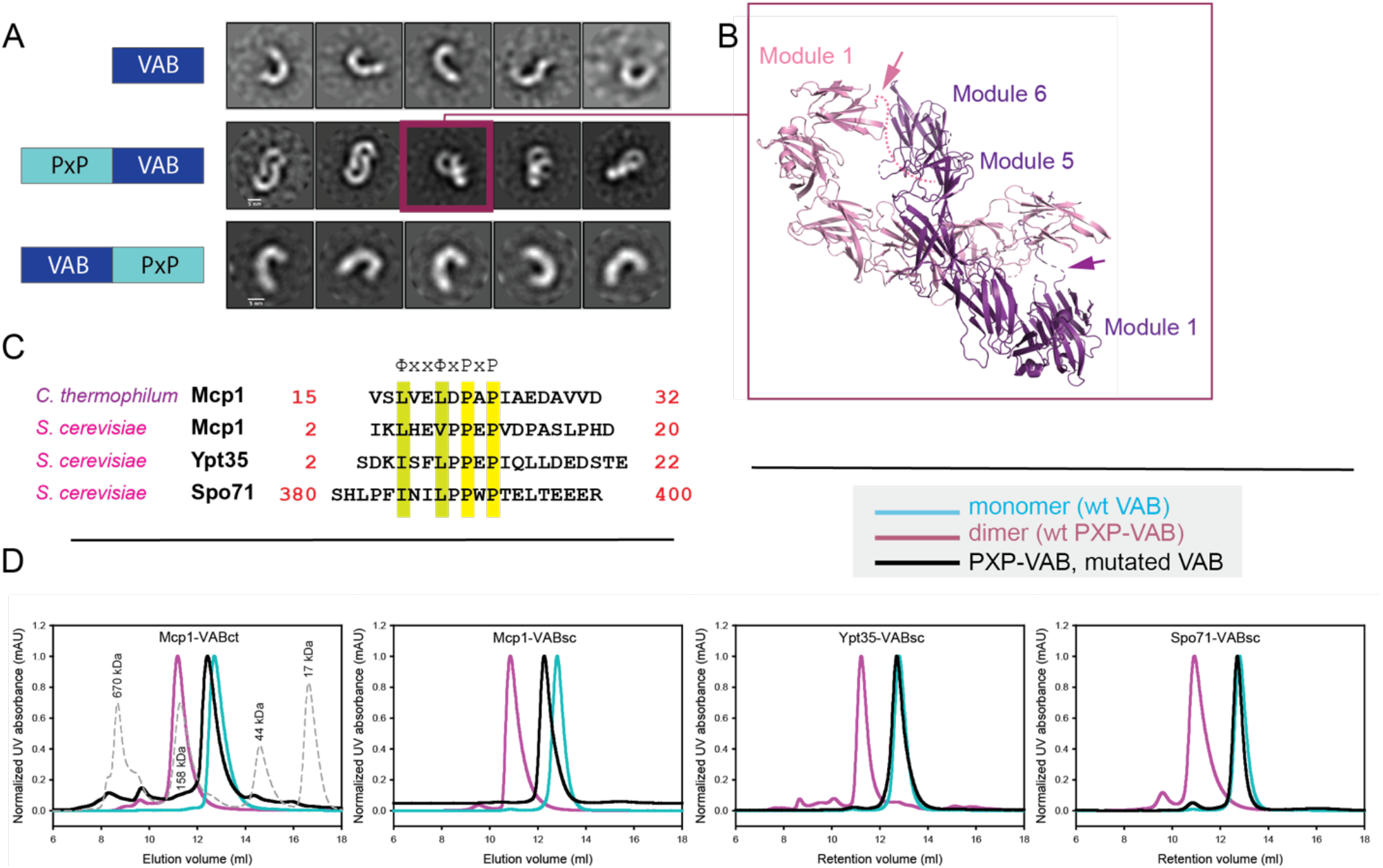
The Pro-X-Pro motifs of Mcp1p, Spo71p, and Ypt35p bind to the same surface of the VAB. (A) The VABct and VAB_*ct*_-PXP(Mcp1_*ct*_) constructs are monomeric in solution as assessed by negative stain electron microscopy, whereas PXP(Mcp1_*ct*_)-VAB_*ct*_ dimerizes. Class averages are shown (scalebar 5 nm). (B) PXP(Mcp1_*ct*_)-VAB_*ct*_ dimerization is in trans, with the N-terminal PXP-motif from one monomer bound to the C-terminal end of the second monomer and vice versa. The dimer in the asymmetric unit of the crystal is similar to the class average boxed in (A). (C) Pro-X-Pro motifs of Mcp1p from *C. thermophilum* and *S. cerevisiae*, and from *S. cerevisiae* Spo71p, and Ytp35p. (D) Size exclusion profiles of wild type and mutant constructs of PXP(Mcp1_*c*t_)-VAB_*ct*_, PXP(Mcp1_*sc*_)-VAB_*sc*_, PXP(Spo71_*sc*_)-VAB_*sc*_, and PXP(Ypt35_*sc*_)-VAB_*sc*_. The wild-type constructs are dimers, indicating an intact Pro-X-Pro binding site. For the mutants, residues important for the binding of the Pro-X-Pro motif as determined from the crystal structure were altered, and the constructs are monomeric. This shows that the Pro-X-Pro motifs of Mcp1p, Spo71p, and Ypt35p all bind this site on the VAB surface.

We used a similar approach to assess whether, as was reported, the Pro-X-Pro-motifs of Spo71p and Ypt35p bind in the same site. We made PXP-VAB_*sc*_ constructs in which we N-terminally fused the Pro-X-Pro-motifs from *S. cerevisiae* Mcp1p (residues 2-20), Spo71p (residues 380-400) or Ypt35p (residues 2-22) (see Figure 2C for sequences) to the *S. cerevisiae* VAB (residues 1869-2545) or a mutated version, in which the residues in the proposed Pro-X-Pro-binding surface were altered as before (R2396E, I2468A, I2473A and M2474A). While the PXP-VAB_*sc*_ constructs with the wild-type VAB sequence dimerized, the mutant PXP-VAB_*sc*_ constructs were monomeric, indicating that the Pro-X-Pro motifs of Mcp1p, Spo71p and Ypt35p all bind in the same site (Figure 2D).

To assess the notion that VPS13 proteins act in collaboration with scramblases, like the VPS13 family protein ATG2, we used a well-established fluorescence-based assay to determine whether multi-span proteins known to interact directly with VPS13 proteins, Mcp1p (Bean et al., 2018; John Peter et al., 2017) and XK (Park and Neiman, 2020), might scramble glycerolipids (Goren et al., 2014; Ploier and Menon, 2016). We overexpressed these proteins, purified them in detergent (n-dodecyl- ß -D-maltopyranoside and glycol-diosgenin for Mcp1p and XK, respectively), and reconstituted them into liposomes containing small (0.5%) amounts of nitrobenzoxadiazole (NBD)-labeled lipid (Figure 3A). The addition of dithionite, a membrane impermeable reducing agent, reduces solvent accessible NBD to quench its fluorescence. In the absence of scramblase activity, dithionite can access NBD only in the outer leaflet of the liposome but not in the liposome lumen, so that dithionite addition results in a ∼50% reduction in fluorescence (Figure 3B). In the presence of a scramblase, which continuously exchanges NBD-lipids between the leaflets of the membrane bilayer, all NBD becomes accessible, resulting in a larger, near complete (>>50%) reduction in fluorescence upon dithionite addition. The reduction would be complete in an ideal reconstitution scenario, where all liposomes incorporate scramblase activity. Using this assay, we found that both Mcp1p and XK scramble glycerolipids non-specifically, including PE, PC, and PS (Figure 3 C-E,G-I), consistent with the ability of Vps13s to transport glycerolipids non-specifically. We ruled out that the observed fluorescence reduction is due to leaky liposomes, for example due to incomplete detergent removal, which would allow dithionite to penetrate into the liposome lumen to reduce NBD there even in the absence of a scramblase (Goren et al., 2014; Ploier and Menon, 2016). For this, we prepared proteoliposomes as before, but lacking NBD-lipid and in the presence of NBD-glucose, and dialyzed the liposomes extensively against NBD-glucose free buffer (Figure 3B). NBD-glucose was retained in the liposomes and was not accessible to dithionite, demonstrating that the liposomes were not leaky (Figure 3F,J). Interestingly, in contrast to XK, the XK-related proteins −8 and-9 are reported to be inactive in vitro in scrambling assays (Sakuragi et al., 2021; Straub et al., 2021), although their scrambling activity was demonstrated in cells (Suzuki et al., 2014). It was suggested that caspase processing or some other still poorly understood post-translational modification is necessary to activate scrambling for these proteins.

**Figure 3.**
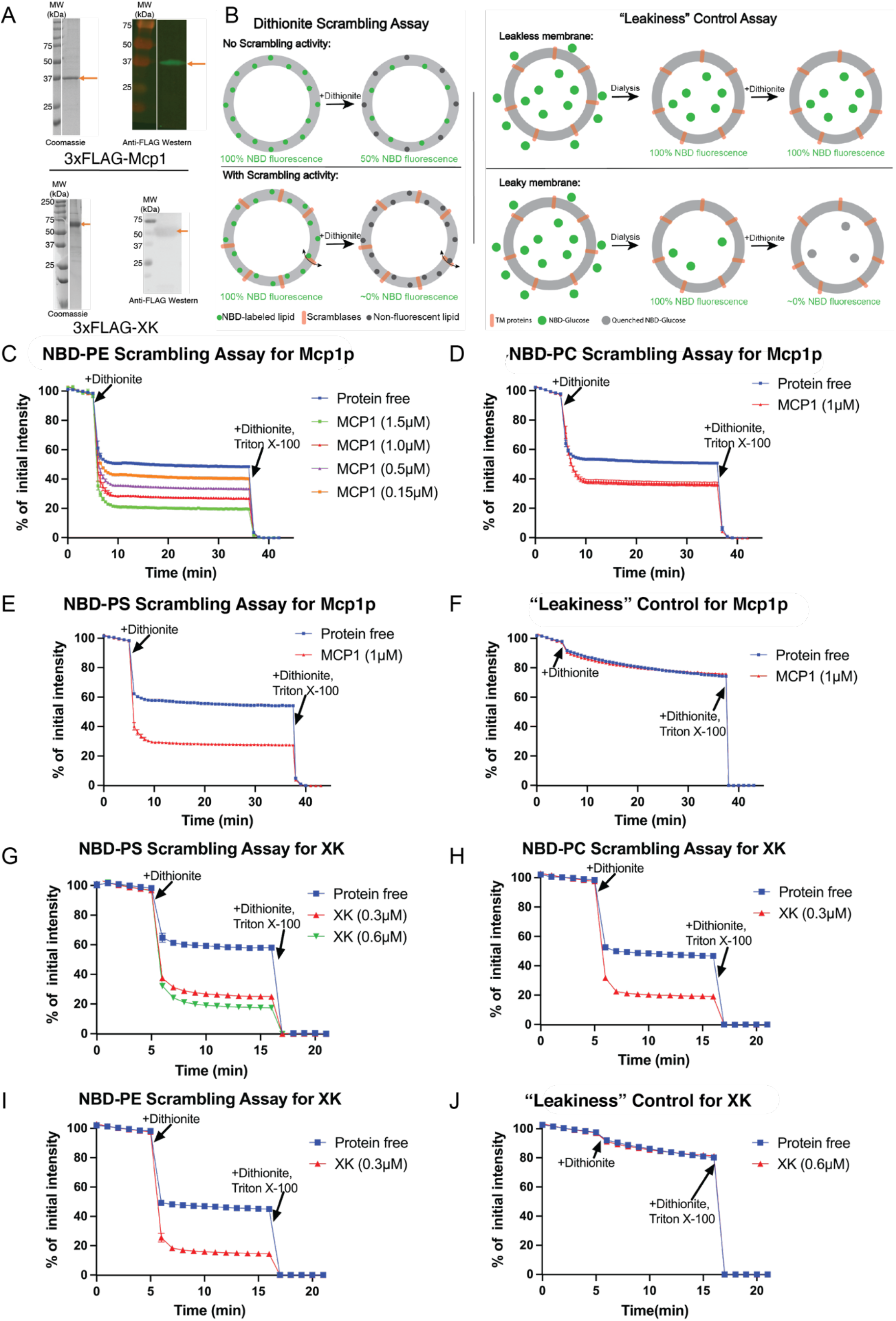
Mcp1p and XK scramble glycerolipids in vitro. (A) SDS-PAGE gels showing purified 3xFLAG-Mcp1p or 3xFLAG-XK before their reconstitution into liposomes, analyzed by Coomassie staining and by Western blotting with anti-FLAG. (B) Schematics for the dithionite scrambling assay and “leakiness” control. (C) Mcp1p scrambles NBD-PE. Scrambling is not observed with the protein free liposomes. Reconstitution is more efficient when the protein is added at higher concentrations, resulting in nearly complete reduction of fluorescence. (D and E) Mcp1p scrambles NBD-PC and NBD-PS. (F) “Leakiness” control for Mcp1p-containing liposomes. Fluorescence retention of NBD-glucose in the liposome lumen after dialysis and, further, after addition of dithionite indicates that the liposome membranes remain intact and impermeable to small molecules like dithionite or NBD-glucose. (G-I) Scrambling results for XK. XK scrambles NBD-lipids without headgroup specificity. (J) The XK-containing liposomes are leak-free.

## Discussion

Our studies of the VPS13 VAB are a step toward understanding how the C-terminal end of VPS13s interacts with membranes. In particular, we have identified the binding site for the Pro-X-Pro motifs of VPS13 receptors on one of the flat faces of the fishhook -shaped VAB, on a highly conserved surface patch covering large portions of the 6^th^ repeat module and including the interface between the 5^th^ and 6^th^ modules. The receptor may make additional interactions with the VAB as the conserved surface is larger than the Pro-X-Pro motif binding site observed in the crystal structure. The large conserved surface centered on the 6^th^ repeat module is also present across metazoan VPS13 VABs and so likely also serves in receptor recognition there.

Noted before, the topology of each VAB repeat module is like that of bacterial lysins. These are proteins that interact with and/or form pores in membranes via loops emanating from one end of the ß-sandwich (for examples, see (Kocar et al., 2021; Yanagihara et al., 2010). In the VAB, these loops are at the interface between modules and so most likely do not play a role in membrane binding. In fact, the VABs from *S. cerevisiae* and *C. thermophilum* VPS13 do not bind membranes in liposome flotation assays, even in the presence of phosphoinositides (8% PI3P, PI4P, or PI(4,5)P_2;_ data not shown). It is plausible, however, that other features at the very C-terminal end of VPS13s, including several amphipathic helices or the PH domain (Fidler et al., 2016; Kumar et al., 2018), might directly bind to membranes, and that the flat partially basic face of the VAB (opposite the Pro-X-Pro binding face) is apposed to membranes when Vps13p is fully assembled with all its interaction partners at contact sites.

Importantly, we have shown that two integral membrane proteins that interact directly with VPS13s, Mcp1p and XK, are scramblases. We and others reported previously that to effect bulk lipid transfer the VPS13 family protein ATG2 interacts with scramblases (Ghanbarpour et al., 2021; Maeda et al., 2020; Matoba et al., 2020) on both donor and acceptor organelles at contact sites; our data here indicate that a collaboration with scramblases is a more general feature of VPS13 family proteins. Mcp1p interacts with Vps13p via the VAB at the C-terminus. Although not required for lipid transport in principle, we speculate that there may be additional interactions between the taco-shell lipid transport domain of VPS13, and in particular the so-called “APT1” region that forms one of its ends close to the VAB, and the transmembrane domain of Mcp1p allowing for a hand-off of lipids between transfer protein, the scramblase in the membrane, and the membrane (Figure 4). Multi-pass integral membrane proteins, including scramblases like Mcp1p, may also accelerate lipid transfer by lowering the rate determining steps, i.e., lipid extraction from or insertion into donor or acceptor membrane, respectively.

**Figure 4.**
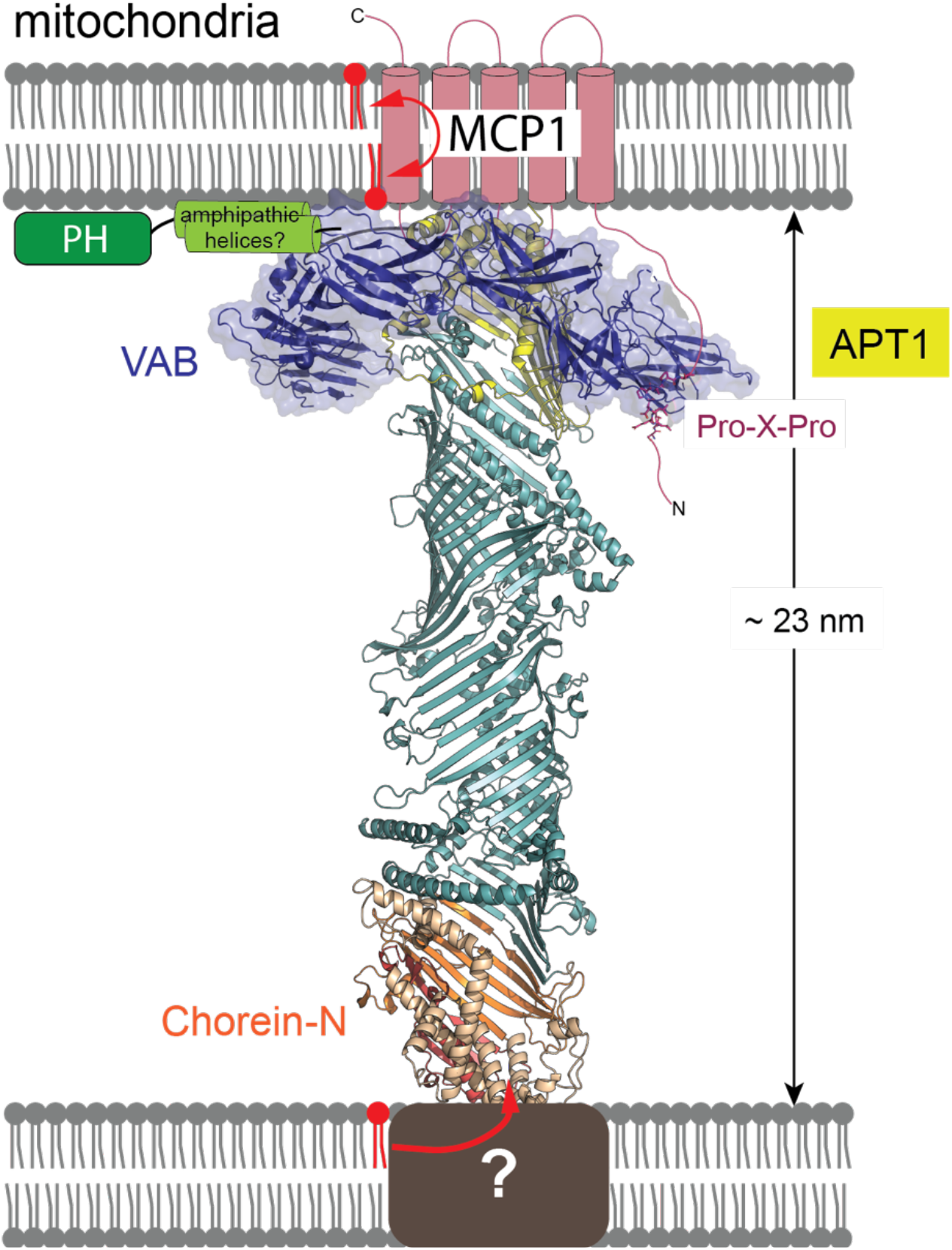
Cartoon of Vps13 bound between membranes and interacting with Mcp1p via its VAB. The “chorein-N” (orange), “extended-chorein” (teal), and APT1 (yellow) domains are believed to form a lipid transport channel. The VAB interacts with the Pro-X-Pro motif in the predicted unstructured N-terminus of Mcp1p (indicated). We speculate that the APT1domain interfaces directly with the integral membrane domain of Mcp1, which harbors scrambling activity, and that there might be a direct hand-off of lipids between Vps13p and the scramblase. The ribbons cartoon of the VPS13 lipid transport module was generated using RoseTTAFold (Baek et al., 2021).

How XK interacts with VPS13A, or which end of VPS13A is involved, is even less well understood, although it is tempting to speculate that in this case also there could be a direct interaction between the lipid transfer domain of VPS13 and the transmembrane domain of XK and a hand-off of lipids. (In this case, the interaction is unlikely to involve the VPS13A VAB, as we were not able to detect in pull down experiments an interaction between XK and the VAB (residues 1877 – 2542 of human VPS13A co-expressed with XK in mammalian cells; data not shown.)

We anticipate that additional scramblases will continue being identified as the known VPS13 interactome expands. It is also possible that the VPS13 proteins also partner with other multi-pass integral membrane proteins that do not have scrambling activity. For example, these could be part of the lipid biosynthetic machinery, allowing lipids to be removed from the membrane leaflet in which they are made to another compartment as they are made. Such a scenario was recently proposed for a newly identified member of the VPS13 family, Csf1p (Toulmay et al., 2022). We predict that this model of a partnership between scramblases, or other classes of multi-pass integral membrane proteins, and lipid transporters will also be applicable to more recently identified members of the VPS13 protein family, including SHIP164, KIAA1109/Csf1p, and Hobbit/Hob1p/Hob2p (Gomes Castro et al., 2021; Hanna et al., 2021; John Peter et al., 2021; Neuman et al., 2022; Toulmay et al., 2022).

Coordinates and structure factors for the VPS13 VAB have been deposited with the PDB (accession no. 7U8T).

## Materials and Methods

### Materials

All the lipids including POPC (Cat.#850457C), POPE (Cat.#850757C), NBD-PE (Cat.#810153C), NBD-PC (Cat.#810122), and NBD-PS (Cat.#810198) were purchased from Avanti Polar Lipids. NBD-glucose was purchased from Abcam (Cat. #186689-07-6) and DDM from GoldBioTech (Cat.#DDM25), GDN from Anatrace (Cat.#GDN101). Bio-BeadsTM SM2 Adsorbent Media was purchased from BIO-RAD (Cat.#152-3920), and anti-FLAG M2 resin was from Sigma Aldrich (Cat.#A2220). EDTA-free Roche cOmplete protease inhibitor cocktail (Cat.#4693159001). Expi293F cells and *E. coli* BL21 (DE3) competent cells were purchased from ThermoFisher (Cat.#’s A14527 and C600003).

## Methods

### Structural and biochemical studies of the VPS13 VAB

#### Plasmid constructs

VAB domain (residues 1944 – 2635) of Vps13p from *C. thermophilum* (Uniprot G0S388) was cloned from genomic DNA into a modified pET-Duet vector with a C-terminal 6xHis tag. To generate PxP(Mcp1ct)-VABct_1944-2635_-His_6_, residues 15 – 32 of Mcp1 from *C. thermophilum* (Uniprot G0S0T5) were fused N-terminally to the VAB_1944-2635_-His_6_ construct. VAB domain (residues 1869 – 2545) of Vps13p from *S. cerevisiae* was similarly cloned from genomic DNA into pET-29a(+) expression vector containing a C-terminal 6xHis tag. PxP(Mcp1sc)- VABsc_1869 – 2545_- His_6_, PxP (Ypt35sc)-VABsc_1869 – 2545_- His_6_, and PxP(Spo71sc)-VABsc_1869 – 2545_- His_6_ constructs contain N-terminal fusion of residues 2 – 20 from yeast Mcp1, residues 2 – 22 from Ypt35 and residues 380 – 400 from Spo71, respectively. All mutants were generated by round-the-horn mutagenesis. See Supplementary Table 1 for a list of primers.

#### Protein expression and purification

All plasmids were transformed in *E. coli* BL21 (DE3) cells. Cells were cultured at 37 °C until O.D. (600 nm) reached 0.8, and protein expression was induced with 0.5 mM IPTG at 18 °C for 16 hours. Harvested cells were resuspended in buffer (20 mM HEPES pH 7.8, 500 mM NaCl, 1 mM TCEP.HCl, 10% glycerol) supplemented with 20 mM Imidazole, 5 mM MgCl_2_, DnaseI and 1x cOmplete EDTA-free protease inhibitor cocktail (Roche), lysed using a cell disruptor (5 passes at 15,000 psi) and clarified by centrifugation at 15,000 rpm for 45 mins. Cell lysate was loaded on Ni^2+^-NTA column pre-equilibrated with lysis buffer. After washing, bound protein was eluted with buffer A containing 500 mM Imidazole, concentrated using an Amicon Ultra centrifugal filter unit with a 50 kDa MWCO, and loaded on a Superdex 200 gel filtration column (GE Healthcare) equilibrated with buffer A.

#### Crystallization, data collection, structure determination

PxP(Mcp1ct)-VABct_1944-2635_-His_6_ was crystallized with sitting-drop vapor diffusion method. Equal volumes of protein at 3 mg/ml concentration and reservoir solution (75 mM Imidazole pH 8.2, 70 mM Li_2_SO_4_, 500 mM NaCl) were mixed and incubated at 295 K. Plate-like crystals, belonging to space group P2_1_, were cryo-protected by serial transfer in mother liquor containing increasing concentrations of glycerol, loop-mounted and flash frozen in liquid nitrogen. Diffraction data was collected at 100 K and a wavelength of 0.979 Å at the NE-CAT beamline 24-ID-E at the Advanced Photon Source, using a Dectris EIGER 16M detector (Dectris Ltd.). All data were indexed, integrated and scaled using XDS (Kabsch, 2010) with the statistics given in Supplementary Table 2. The structure of PxP(Mcp1ct)- VABct_1944-2635_-His_6_ was solved by molecular replacement with Phaser MR (McCoy et al., 2007), using a model generated with the AlphaFold2 Colab notebook (Jumper et al., 2021) pre-processed with phenix.process_predicted_model (Liebschner et al., 2019) to remove low confidence residues with pLDDT scores < 70. The first three repeat modules were initially searched and confidently placed in the electron density, followed by the next two modules in both chains. Finally, the sixth repeat module was identified only for one of the chains (PxP-VAB_1-6_). Following jelly body refinement, strong and continuous density corresponding to the Pro-X-Pro peptide appeared in 2Fo-Fc maps (Figure 1D). In modeling the peptide into this density, we were guided by the restraints imposed by the linker length, as the Pro-X-Pro peptide binds in trans, stretching from the N-terminus of one molecule to its binding site on the other. This restraint set the directionality of the peptide chain. A prominent feature of the backbone density was a kink, into which the Pro-Ala-Pro sequence was modeled, with both prolines featuring cis peptide bonds. Maps calculated once these residues and the rest of the peptide’s polyalanine backbone had been placed showed good side chain density, allowing us also to model side chains for other residues in the Pro-X-Pro motif. We further checked difference maps to adjust side chain positions during refinement. The peptide register as modeled was the best fit to this density. The refinement consisted of cycles of manual model rebuilding in COOT (Emsley et al., 2010), individual isotropic b-factor and TLS refinement with REFMAC5 (Murshudov et al., 2011) and phenix.refine (Liebschner et al., 2019). Refinement statistics are given in Supplementary Table 2.

Structures were rendered in PyMol (The PyMOL Molecular Graphics System, Version 2.0, Schrödinger, LLC).

### Scrambling Studies for Mcp1p and XK

#### Plasmids

The coding sequence of MCP1 (Uniprot Q12106) and XK (Uniprot P51811) were PCR amplified from the *S. cerevisiae* genomic DNA and human cDNA library, respectively, and subcloned into pCMV10 vector with an N-terminal 3xFLAG tag and PreScission protease cleavage site.

#### Membrane protein expression and purification

Constructs encoding MCP1 or XK were transfected into Expi293F cells (Thermo Fisher Scientific) according to manufacturer’s instructions. The cells were harvested 48 hours after transfection.

For Mcp1p, the cells were pelleted, and resuspended in buffer B (50 mM HEPES, pH7.0, 500 mM NaCl, 1 mM TCEP, 10% glycerol) containing 1× cOmplete EDTA-free protease inhibitor cocktail (Roche) and lysed using a Dounce homogenizer (15∼20 passes). To solubilize the protein, powdered DDM was added to the lysate at a final concentration of 1% w/v, and the lysate was gently agitated in the cold room for 90 minutes. Cell lysates were clarified via centrifugation at 100,000 *g* for 60 minutes and supernatant was incubated with anti-FLAG M2 resin (Sigma Aldrich), which was pre-equilibrated with buffer C (buffer B, 0.02% DDM), at 4°C for 2 hours, and then the resin was washed with buffer C. To remove chaperone, resin was incubated with buffer B containing 5 mM MgCl_2_, 2.5 mM ATP containing buffer C at 4 °C overnight. Bound proteins were further washed with buffer C, then eluted using 0.25mg/ml 3xFlag peptide in Buffer C. The proteins were concentrated in a 10-kD molecular weight cutoff (MWCO) Amicon centrifugal filtration device (UFC501024) and quantified by Coomassie blue staining using BSA standards.

XK was purified similarly to Mcp1p but with buffer D (50 mM HEPES, pH 7.8, 200 mM NaCl, 1 mM TCEP, 10% Glycerol) and with GDN instead of DDM throughout (1.5% GDN for solubilization, 0.02% GDN for purification). XK was further gel filtrated with Superdex 200 increase 10/300 column (Cytiva).

#### Liposome preparation

90% POPC, 9.5% POPE, and 0.5% NBD-labeled lipid (NBD-PE, NBD-PC, or NBD-PS) in chloroform were mixed and dried under a N_2_ stream, and further vacuum dried for 30 minutes. The resulting lipid film was resuspended in buffer E (50 mM HEPES, pH 7.0, 200 mM NaCl) for MCP1 and in buffer F (50 mM HEPES, pH 8.0, 200 mM NaCl) for XK to generate a 10.5 mM lipid stock. The mixture was incubated at 37°C for 60 minutes, with vortexing every 10 minutes, and then freeze-thawed for ten cycles. Liposomes were extruded 31 times through a 400 nm polycarbonate filter and used within 6 hours.

#### Proteoliposome preparation

Proteoliposomes were prepared as described in (Marek and Gunther-Pomorski, 2016; Ploier and Menon, 2016). Briefly, liposomes at final lipid concentration of 5.25 mM and in 250 μl total volume were destabilized by addition of Triton X-100 to a final concentration determined by the swelling titration assay as described in (Ploier and Menon, 2016). The final Triton X-100 concentration was 3 mM. After 2 hours of destabilization at room temperature, detergent solubilized proteins (MCP1 or XK) were added and the samples were gently agitated for an hour to allow protein incorporation. The detergent was removed in three steps using pre-washed biobeads: following a first addition of biobeads (20 mg), the sample was incubated at room temperature for an hour; then another aliquot of biobeads (20 mg) was added and the sample was rotated at room temperature for two more hours. Finally, the sample was pipetted into a new tube containing fresh biobeads (40 mg) and agitated at 4°C overnight. Biobeads were removed by pipetting, and the sample was dialyzed against buffer E for MCP1 and buffer F for XK for two days at 4°C. The protein reconstitution efficiency was ∼100% as assessed using a liposome flotation assay (Karatekin and Rothman, 2012).

#### Scrambling assay

The scrambling assay (Goren et al., 2014; Ploier and Menon, 2016) was performed at 30°C in 96-well plates, with 100 μl reaction volumes of liposomes/proteoliposomes (∼260 μM final lipid concentration) prepared as described above. To assess scrambling, NBD fluorescence after addition of dithionite (to 5 mM) was monitored (excitation at 460 nm, emission at 538 nm) using the Synergy H1 Hybrid Multi-Mode Reader (BioTek). Finally, additional diothionite (5 mM) and Triton X-100 (0.5%) were added. The Triton X-100 dissolves the liposomes, allowing complete quenching of the NBD.

A similar protocol was used for the NBD-glucose leakiness control assay (Goren et al., 2014; Ploier and Menon, 2016), except that no NBD-lipids were incorporated into the liposomes or proteoliposomes. Instead, NBD-glucose (3 mM) was added during the destabilization step.

## Acknowledgements

We thank T. Melia and P. De Camilli for critically reading this manuscript and the staff of the Northeastern Collaborative Access Team (NE-CAT) beamlines of the Advanced Photon Source (Argonne National Laboratory) for their help with data collection. NE-CAT is funded by National Institute of General Medical Sciences of the National Institutes of Health (NIH) (P30 GM124165), NIH-ORIP HEI grant (S10OD021527), and contract DE-AC02-06CH11357. This research was funded by grants from the NIH (R35GM131715) and by Aligning Science Across Parkinson’s grant ASAP-000580 through the Michael J. Fox Foundation for Parkinson’s Research (MJFF) to KMR. ZH was supported by the China Scholarship Council. For the purpose of open access, the author has applied a CC BY public copyright license to all Author Accepted Manuscripts arising from this submission. The authors declare no competing financial interests.

## Contributions

KMR conceived of and supervised experiments. PL designed the VAB constructs used in crystallization and obtained initial crystallization conditions. JA optimized crystallization and cryo-conditions, determined and analyzed the structure, and carried out all VAB-related biochemistry. ZH made proteins for and carried out all scrambling assays. JA and ZH made the figures. KMR wrote the manuscript together with JA and ZH.

