## Supplemental Figure 1, Table 1, Table 2 for "Structural and biochemical insights into lipid transport by VPS13 proteins"

**Supplementary materials**:

Supplementary Table 1. Primers for the study.

| | | |
|---|---|---|
| VABct | FP | actttaagaaggagatataccatggtagcgccatacaggatcagg |
| | RP | gttcgacttaagcattatgcggccgcttattagtgatggtgatggtgatgctggcggtacaggctctttg |
| PXP(Mcp1ct)_VABct | FP | catcgccgaggatgccgtcgtcgacgcgccatacaggatcagg |
| | RP | ggggcaggatccaactcgaccagcgataccatggtatatctccttcttaaag |
| VABct_PxP(Mcp1ct) | FP | gtcgagttggatcctgcccccatccatcaccatcaccatcactaataag |
| | RP | cagcgagacggccgagacgagcgtctggcggtacaggctctt |
| VABct_R2489E | FP | gaacagcggctgatccgcgtggag |
| | RP | ctgcccagcgcgagcaatttttag |
| VABct_mutLIM | FP | ccgcgtagtgcagcgccgtatgcctgggacttcccggctgcgaag |
| | RP | gggcgcacggtaccggacaggacgccaaccggagcggtc |
| VABsc | FP | tttaagaaggagatatacatatgaagccatatcaactggtaaac |
| | RP | tggtggtggtggtgctcgagattggccttatagttaacaataactaag |
| PxP(Mcp1sc)-VABsc | FP | agtagaccctgctagtctccctcatgataagccatatcaactggtaaac |
| | RP | ggttctggaggcacttcatgcaactttatcatatgtatatctccttcttaaag |
| PxP(Ypt35sc)-VABsc | FP | catccaactacttgacgaagactccacggagaagccatatcaactggtaaac |
| | RP | ggttcgggaggtagaaaggatatcttgtcgctcatatgtatatctccttcttaaag |
| PxP(Spo71sc)-VABsc | FP | atggcctaccgaactgacggaggaagagagaaagccatatcaactggtaaac |
| | RP | ggaggtaggatattaataaacggaagatgactcatatgtatatctccttcttaaag |
| VABsc_R2396E | FP | gaacataagcttttaaagattgaaattcttttggacaaagc |
| | RP | tgaatttttcaaaactttcaaatatgttacaccgacatttttg |
| VABsc_mutIIM | FP | tcgaaaagtgcggcgccatacgcatgggattttcctacagctaaggag |
| | RP | gggcgccctgtaaaagataggtttgaaactacggcttga |
| Mcp1sc | FP | aggggccccttgcggccgccatgataaagttgcatgaagtgcc |
| | RP | agggatgccacccgggatccctaattcacgtgcaacagc |
| XK | FP | ccaggggccccttgcggccgcgatgaaattcccggcctcg |
| | RP | agggatgccacccgggatccttaagcagagcagagatcttc |

Supplementary Table 2: Data collection and refinement statistics.

| | |
|---|---|
| **Crystal parameters** | |
| Space group | P 1 21 1 |
| Cell dimensions | |
| a, b, c (Å) | 83.735, 91.225, 132.769 |
| α, β, γ (°) | 90.000, 102.163, 90.000 |
| Monomers/ASU | 2 |
| **Data collection** | |
| Wavelength (Å) | 0.979180 |
| Resolution range (Å) | 48.38-3.00 (3.11-3.00) |
| Completeness (%) | 98.5 (97.6) |
| Redundancy | 3.5 (3.6) |
| I/σ(I) | 17.2 (3.4) |
| Rmerge (%) | 4.4 (32.2) |
| Rmeas (%) | 5.2 (37.8) |
| Rpim (%) | 2.8 (19.7) |
| CC (1/2) | 0.999 (0.970) |
| Refinement | |
| Resolution range (Å) | 48.36-3.00 (3.07-3.00) |
| Rwork (%) | 24.96 |
| Rfree (%) | 28.48 |
| RMSD bond angles/lengths | 0.852/0.005 |
| Ramachandran statistics (% in favored /allowed/other regions) | 95.96/3.86/0.18 |
| PDB accession code | 7U8T |

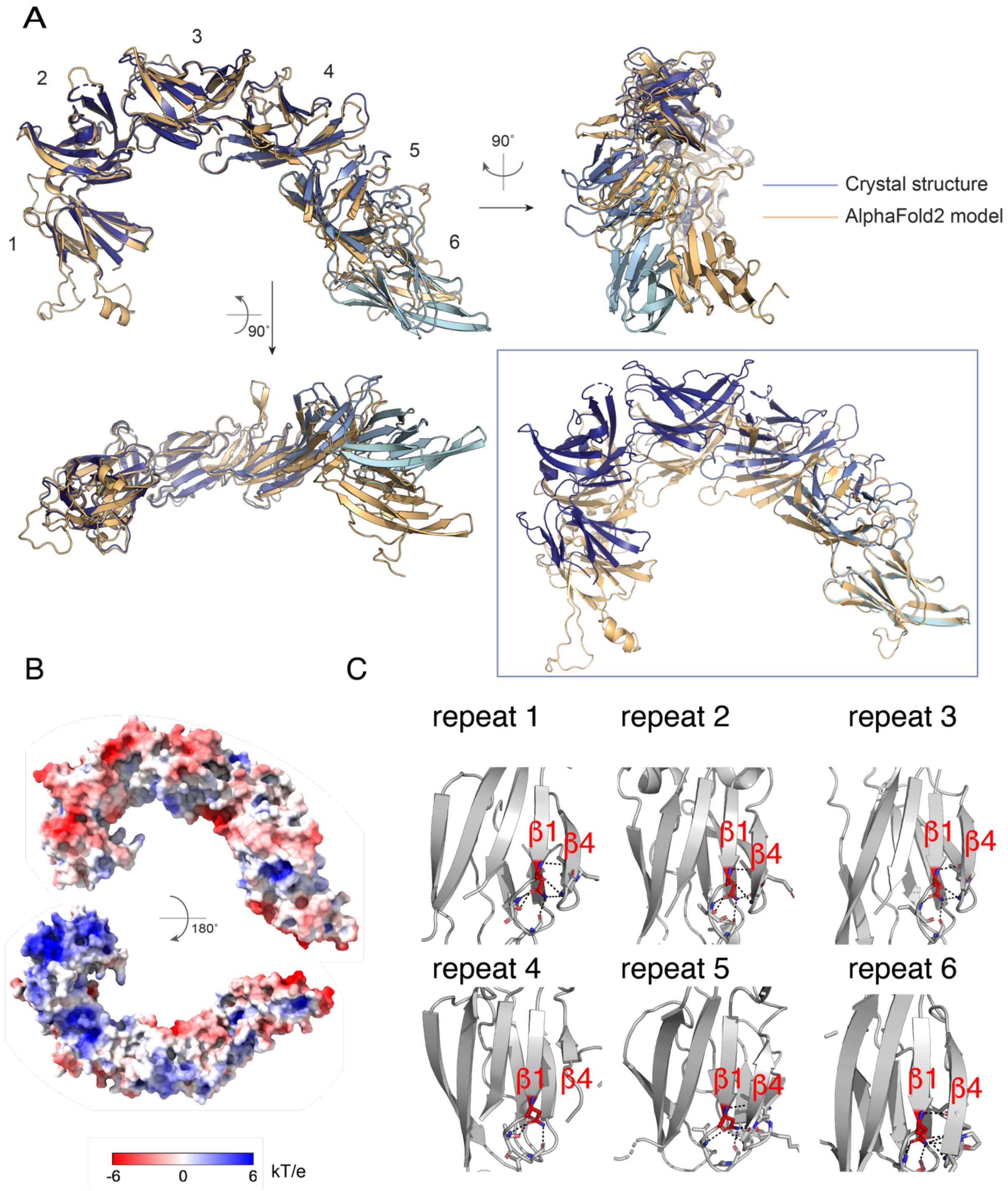


Supplementary Figure 1.

Supplementary Figure 1. **Details of the VAB structure.** (A) Comparison of PXP-VAB$_{1-6}$ in the crystal structure with the prediction from AlphaFold2. AlphaFold2 accurately predicted the fold of the individual modules as well as the interfaces between modules 1&2 and 2&3. The remaining interfaces in the crystal structure differed from those in the AlphaFold2 model. At left, the structures are superimposed based on the first three modules. The positions/orientations of the remaining modules differ in the two structures. In the box, the structures are superimposed based on the 6$^{th}$ module only. The interface between modules 5&6, where the Pro-X-Pro motif binds, is different in the two models. The AlphaFold2 model does not feature the groove that is the Pro-X-Pro motif binding site. (B) The electrostatic potential as calculated by APBS software (Jurrus et al., 2018) mapped onto the surface of the VAB. The VAB is shown in the same orientations as in Figure 1C. (C) The asparagine at the end of ß-strand 1, strictly conserved in all repeat modules, is involved in an extensive hydrogen bonding network that stabilizes folding of the module.

Jurrus, E., D. Engel, K. Star, K. Monson, J. Brandi, L.E. Felberg, D.H. Brookes, L. Wilson, J. Chen, K. Liles, M. Chun, P. Li, D.W. Gohara, T. Dolinsky, R. Konecny, D.R. Koes, J.E. Nielsen, T. Head-Gordon, W. Geng, R. Krasny, G.W. Wei, M.J. Holst, J.A. McCammon, and N.A. Baker. 2018. Improvements to the APBS biomolecular solvation software suite. *Protein Sci*. 27:112-128.
